# 3D printed microtissue cassettes enabling high throughput proton radiobiological assays

**DOI:** 10.1101/2024.08.10.607473

**Authors:** Chih-Tsung Yang, I-Chun Cho, Ching-Fang Yu, Edward Cheah, Tesi Liu, Yi-Ping Lin, Sing-Yu Hu, Jyun-Wei Jheng, Ivan Kempson, Tsi-Chian Chao, Sen-Hao Lee, Eva Bezak, Benjamin Thierry

**Affiliations:** Future Industries Institute, University of South Australia, Adelaide, Australia; Research Center for Radiation Medicine, Chang Gung University, Taoyuan, Taiwan; Radiation Research Core Laboratory, Linkou Chang Gung Memorial Hospital, Taoyuan, Taiwan; Department of Radiation Oncology, Chang Gung Memorial Hospital Linkou Branch, Taoyuan, Taiwan; Department of Medical Imaging and Radiological Sciences, Chang Gung University, Taoyuan, Taiwan; Allied Health and Human Performance Academic Unit, University of South Australia, Adelaide, Australia

## Abstract

Uncertainties on proton relative biological effectiveness (RBE) across the spread out of Bragg peak (SOBP) (typically assumed to be 1.1) may lead to suboptimal treatment plan and unwarranted toxicity to organs-at-risk. Herein, we report a reliable analytical method to determine the proton RBE along the SOBP and distal fall-off region. The 3D microtissue cassette enables the high throughput assessment of biological assays including clonogenic assay and γ-H2AX assay following a single proton irradiation. Clonogenic assay shows the RBE of 1.6 (10% cellular survival) which is consistent with the deter-mined RBE of 1.58 using the γ-H2AX assay. Besides, we also show that the high spatial resolution of the cassette can distinguish the minute but significant foci changes (number, area) in response to small proton radiation dose fraction. The results validate the reliability of our setup in addressing critical proton radiobiological questions.

## Introduction

The presence of the Bragg peak in proton beam and the resulting ability to precisely deposit the radiation dose within the tumor volume physical, is driving its increasing used in contemporary radiotherapy for cancer management. In the clinical radiotherapy settings, the proton beam is usually delivered through the spread out of Bragg peak (SOBP) to enhance the proton interrogation of the tumor tissue while mitigating the dose delivered to healthy tissue. Evidence suggests that proton have distinctive radiobiological effects compared to X-ray used in standard radiotherapy. When designing treatment plans, the relative biological effective-ness (RBE) is used to convert the absorbed dose to describe a response equivalent to photon treatments. Current proton treatment plan usually relies on the historical assumption that the generic value of proton RBE is 1.1. However, it is now well acknowledged that the RBE varies substantially based on both physical factors (dose, dose rate, linear energy transfer, fractionation) and biological ones.^1^ For example, the RBE has been shown to increase with decreasing energy and dose, both of which varies substantially within the treated gross tumor volume. Experimentally, significant variations of the RBE across the SOBP have been reported (e.g. ∼1.1 at the entrance, ∼1.15 at the centre, ∼1.35 at the distal edge and ∼1.7 at the distal fall-off).^2^ A recent review reports that the maximum target RBEs based on clinical and in vitro cell survival data across various cancers are 1.7 and 2.3, respectively.^3^ The exponential increase of the linear energy transfer of pro-ton beams at the distal fall-off of the SOBP has substantial impact on the ambiguity of the RBE. For instance, a RBE of 3.5 was calculated based on U87 cells and the RBE variation effect was more evident for radiosensitive cells at the distal fall-off..^4^ For example, HTB 140 melanoma cells were irradiated with proton at 2 Gy and the RBE was determined to be 7.14 at the distal fall-off.^5^ Quantification of γ-H2AX foci associated to the repair of DNA double strand breaks revealed that the RBE at 6 mm beyond the Bragg peak is 4 (3 h post irradiation) and significantly increased to 6 at 24 h post proton irradiation.^6^ These uncertainties on proton beam RBEs in the clinical settings and more generally on the radiobiological effects may lead to suboptimal local control and unwarranted toxicity to organs-at-risk. This knowledge gap emphasizes the critical need for more radiobiological research and particularly for a robust experimental and standardized method to quantify the correlation between the RBE and biological readouts.

Several in vitro setups have been proposed to address the ambiguity of proton RBE. For instance, a jig was designed to spatially map the biological effectiveness associated with the linear energy transfer (LET) based on the clonogenic assay.^7^ However, due to the inherent exponential nature of LET spanning from the entrance to the peak of SOBP, this setup fails to uniformly sample the biological effectiveness along the proton SOBP, particularly in the distal fall-off region. In addition, using a 96-well plate increases the throughput, while the maximum number of seeding cells limits the feasibility of assessing high survival fraction based on clonogenic assays. A parallel plate chamber was designed to hold a microscopic slide with the support of worm drive scanner and position the sample to be interrogated with proton beam at 1-mm increment across the SOBP. *In vitro* γ-H2AX assay was used to measure the biological endpoint of DNA double strand damages and repairs to further quantify the RBE. However, this low-throughput setup can only accommodate one sample at a specific position for proton irradiation each time which substantially increases the burden for personnel and difficult to access clinical facilities.^6^

In this work, we aim to provide a reliable analytical approach to address the long-standing radiobiological question associated to proton RBE and more generally charged particle radiation therapy. To this end, we use 3D printing technology to fabricate a microtissue cassette with high spatial resolution which enable rigorous measurement of proton radiobiological effects versus the proton energy along the SOBP in a high throughput manner. The state-of-the-art 3D printer allows printing resolution down to sub-millimeter scale enabling the bandwidth of the microtissue samples to be precisely interrogated along the SOBP. We first validate the reliability of the microtissue cassette using the gold standard clonogenic assay to determine the cell survival fraction and RBEs along the proton SOBP, followed by analysis of DNA DSBs repairs including γ-H2AX fluorescence quantification (to determine biological RBE), foci numbers and morphologies.

## MATERIALS AND METHODS

### Materials

U87 glioblastoma cells were purchased from ATCC. High glucose Dulbecco’s Modified Eagle Medium was obtained from GibcoTM. iBidi-treat plastic cover slips (7.5 cm x 2.5 cm) were purchased from iBidi. Alexa Fluor® 488 anti-H2A.X Antibody was purchased from Biolegend. Phospho-Histone H2A.X (Ser139) Antibody #2577 was supplied by Cell Signaling Technology.

### Methods

#### Fabrication of microtissue cassette

The 3D drawing of the microtissue cassette was designed using Autodesk Illustrator and 3D printing was carried out with a Stratasys J735 printer using the VeroClear resin at the printing resolution of high mix 27-micron layer height, followed by water cleaning. The design comprises 16 positions with 0.3 mm in width to accommodate plastic cover slips and 1 mm railings to spatially separate the slides.

#### Preparation of PDMS wells

Polydimethylsiloxane (PDMS) block was prepared by casting 50 mL of PDMS (10:1) in a 10 mm petri dish cured at 65 °C for 6 h. The square PDMS wells was made by cutting an area of 18 mm x 18 mm out of 22 mm x 22 mm PDMS blocks using a scalpel. 2 mm of width on both sides of the square well fits the pillar width in the cassette, providing a surface area of 18 mm x 18 mm for cell seeding.

#### Preparation of microtissue slides for irradiation

U87 cells were cultured in a 10 mm petri dish in an incubator supplied with 5% of CO_2_ at 37 °C. Prior to the cell seeding, the square PDMS well was pressed against the iBidi-treat plastic cover slip. Cells were trypsinized and various cell numbers depending on the biological assays were seeded in the PDMS well, followed by transferring into a 6 well-plate in the incubator for 24 h. One hour prior to irradiation, the PDMS wells were detached from the slides, which were then by transferred to specific positions at 3 mm intervals (to be matched with the water equivalent depth of SOBP) in the microtissue cassette. The lid was closed, and the cell culture medium was aspirated through the opening in the lid and bubbles were removed from the cassette. (Note: Bubbles entrapped in the cassette would affect the proton irradiation). A small piece of PDMS was used to seal the hole and the cassette was gently wrapped with parafilm for irradiation.

### Irradiation setup

The proton irradiation experiments were conducted at the Radiation Research Core Laboratory of the Linkou Chang Gung Memorial Hospital Proton Center. Photon irradiation was performed using the LINAC X-ray at the Department of Radiation Oncology, Chang Gung Memorial Hospital. Following irradiation, samples were maintained and analyzed in the Core Laboratory of Radiation Oncology at CGMH.

For proton irradiation, the cell samples were irradiated with 230 MeV protons on a horizontal beam line. A rotational wheel energy degrader was installed on the proton beam line to create a 6 cm wide SOBP. The dose variation distribution across the SOBP region was within 5%. To eliminate the influence of the proton dose rate on the radiobiological response, the proton dose rate was set at 2 Gy/min, matching clinical settings. The irradiation dose ranged from 2 to 10 Gy. During irradiation, three microtissue cassettes were placed in a customized HDPE cassette holder and precisely positioned for proton irradiation, as shown in the inset image of Figure 1C.

**Figure 1.**
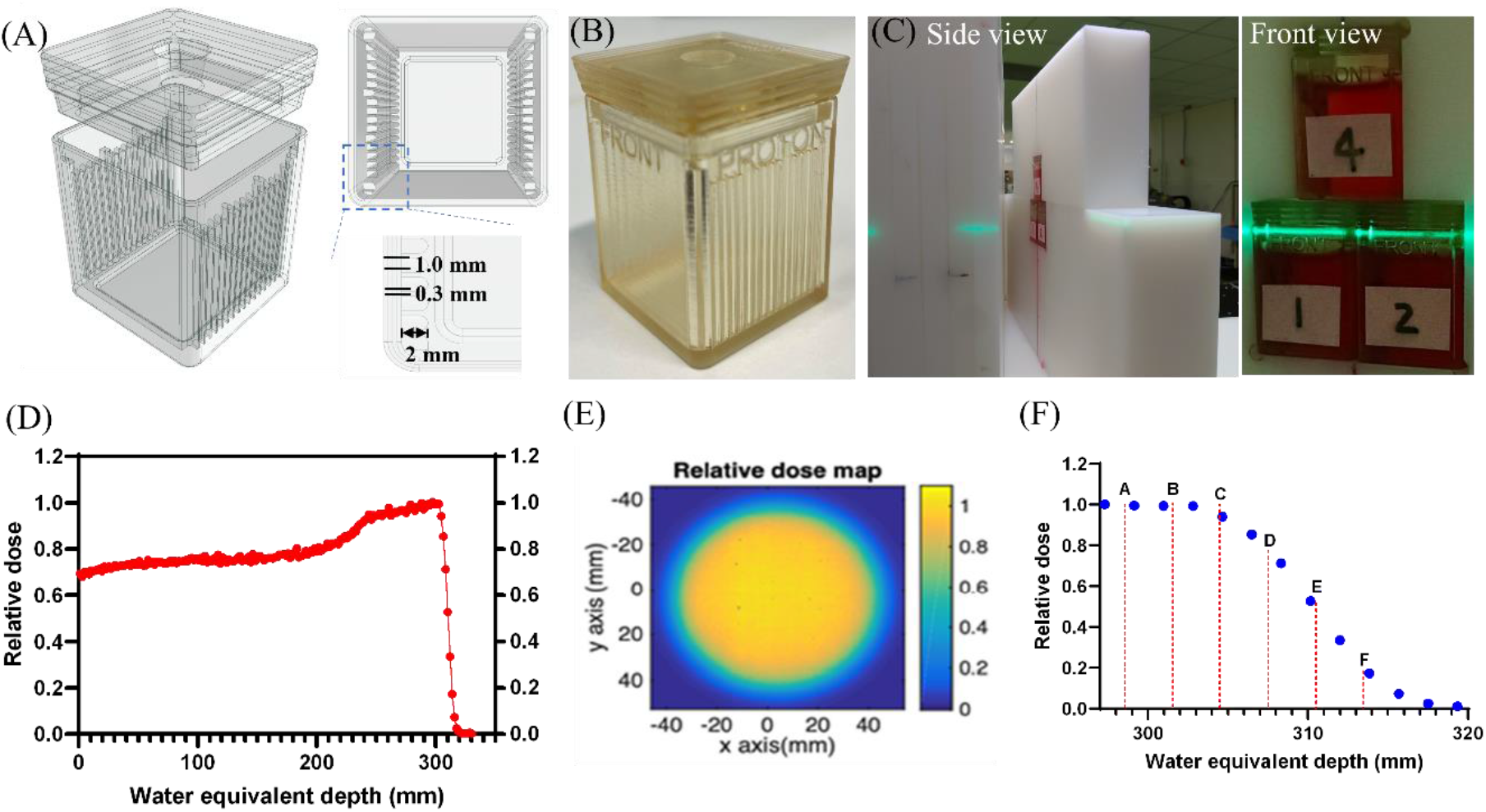
(A) 3D design and geometry of the microtissue cassette; (B) Photograph of a 3D printed microtissue cassette; (C) Three microtissue cassettes in the HDPE block; (D) Profile of the proton beam line at CGMH; (E) The relative dose map of the proton beam line; (F) Correspondence for the 6 slides positions (A to E) in the microtissue with water equivalent depth along the SOBP.

For photon irradiation, cells grown on the iBidi-treat slides and placed in a microtissue cassette were irradiated using a Varian Clinac iX linear accelerator with a 6 MV photon beam. The dose rate was set at 400 Monitor Units per mi-nute (MU/min), equivalent to 4 Gy/min. The dose variation distribution across the microtissue cassette was negligible (∼3%), and three plastic cover slides were placed in the middle position inside the cassette for irradiation.

After irradiation, the cassettes were removed from the HDPE block and transferred to a lead radiation shielding platform for further operations. At the shielding platform, the irradiated slides were transferred to a cassette containing fresh medium to prevent the cells from being affected by oxidative species in the medium. Subsequently, the slides were retrieved and placed in a six-well plate for further culture in the incubator. The slides were analyzed based on the end-point assays.

### Flow cytometry

U87 cells (6 × 10^4^) were seeded in the PDMS well as de-scribed above, and then transferred into the microtissue cassette for photon and proton irradiation at 1 Gy. Following irradiation, cells were trypsinized and incubated with 200 uL of fixation buffer at 4oC overnight. 400 uL of permeabilization buffer was added into the tube, followed by centrifugation at 1500 g for 5 min. After removing the supernatant, 200 uL solution containing 1 uL Fc block and 5 uL goat serum blocking buffer (1% BSA + 0.1% Triton-100X in 1× PBS) was added into the tube and incubated for 30 min at RT. 400 uL of permeabilization buffer was added into the tube, followed by centrifugation at 1500 g for 5 min. The supernatant was removed, followed by the addition of 200 uL of Alex Fluor® 488 anti-H2A.X antibody/isotope (1:200 dilution in 1× permeabilization buffer) into the tube which was kept in the dark for 30 min. 400 uL of 1× permeabilization buffer was added into the tube which was then centrifuged at 1500 g for 5 min, followed by removal of the supernatant. After adding 200 uL of 1× PBS, the solution was analyzed by flow cytometry (BD LSRFortessa™ Cell Analyzer and FlowJo (Tree Star, Inc.) software.

### High content imaging

U87 cells (4 × 10^4^) were seeded on iBidi cover slips as described above and transferred to the microtissue cassette to receive the prescribed proton/photon dose. Cells were fixed and then immunostained with Phospho-Histone H2A.X (Ser139) antibodies at 1h after irradiation. Samples were analyzed by ImageXpress® Micro Confocal at 40X magnification with the exposure parameters for DAPI and FITC at 100 ms and 150 ms, respectively. Image analysis was performed by MetaXpress software to quantify the foci numbers and areas.

### Clonogenic assay

Clonogenic assays were performed to measure cell survival after photon and proton irradiation. 10 × 10^3^ U87 cells were culture on iBi-treated plastic coverslip for 24h before irradiation with photon (0, 1, 2, 4, 8, 10 Gy) and 4 Gy for proton (at SOBP distal fall-off in water: 298.5, 301.5, 304.5, 307.5, 310.5 and 313.5 mm). Clonogenic assays were carried out in triplicates. Following irradiation, various cell number (pho-ton: 0 Gy-400 cells, 1 Gy-1200 cells, 2 Gy-1200 cells, 4 Gy-800 cells, 8 Gy-2400 cells, 10 Gy-2400 cells; proton: 800 cells for each position at the SOBP and distal fall-off) were plated in a 10-mm petri dishes and incubated for one week to allow for the formation of colonies, which was followed by fixation, staining, and manual counting. A colony was determined by counting > 50 cells present in the group.

### Statistical analysis

All experiments were performed independently three times. Experimental data was analyzed with Prism 9 software. t-tests analysis of variance was conducted for statistical analysis. P-values of <0.05 were considered as statistically significant. * p < 0.05, ** p < 0.01, **** p < 0.0001, ns = not significant.

## RESULTS AND DISCUSSION

### Fabrication of 3D printed microtissue cassette and proton irradiation setup

The microtissue cassette was designed by Autodesk Illustrator and contains railings with 1 mm spacing used to position the plastic coverslips as well as an opening at the center of the lid used to remove air bubbles in the cell culture medium (Fig. 1A). Of note, the printer is capable of printing sub-millimeter scale of spacing, which could readily provide higher spatial resolution to investigate proton SOBP with higher spatial accuracy if needed. In this work, the pillar spacing was set at 1 mm and 0.3 mm for the inter-pillar spacing used to position the plastic cover slip. The microtis-sue cassettes (Fig. 1B) were 3D printed using a biocompatible resin. For the proton irradiation setup (Fig. 1C), three microtissue cassettes can be placed simultaneously in the HDPE holder for irradiation to increase the throughput. The clinical proton beam profile used in this study is shown in Fig. 1D and is characterized by a SOBP width R_95p_-R_95d_ of 6.45 cm and a 6 cm of proton field size (Fig. 1E). Fig. 1F illustrates the 6 positions used in the study including 3 in the SOBP and 3 at the SOBP distal fall-off along the SOBP. Prior to irradiation experiments, U87 cells were seeded on iBidi-treated plastic cover slides at a density of 50,000 for 4h. The slides were then transferred into a microtissues cassette to assess the biocompatibility of the 3D printed cassettes. Cells were in good condition as evidenced by cell numbers and morphologies as compared with that in 6-well plate after 48h of incubation.

### Determination of proton RBE based on clonogenic assay

The gold standard clonogenic assay was first used to determine the proton RBE. The clonogenic assay data for photon and proton irradiation are shown in Fig. 2A. and B. For photon irradiation, the colony number was found to correlate as expected with the radiation doses ranging from 0 to 10 Gy. For proton irradiation, a dose of 4 Gy was deposited on plastic slides positioned at six positions along the SOBP. Clonogenic assay results correlated with the PDD of each position along the SOBP. Cell survival at each dose point and position along the SOBP was calculated based on the plating efficiency of the control (0 Gy). The standard linear quadratic model used to fit the data is defined as:

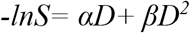

where S is the surviving fraction for a given dose D, and α and β are fit parameters determined by fitting the model to the cell survival data. RBE values at 90%, 50% and 10% of cell survival were calculated to be 1.3, 1.5 and 1.6, respectively. These RBE values are consistent with previous re-ports that the RBE is substantially higher than 1.1 at the SOBP distal fall-off.^2^ Importantly, the proposed methodology enables to perform with high spatial resolution along the SOBP clonogenic as-says in a single irradiation, which may be anticipated to reduce the burden of conducting research proton beam irradiation on technical personnel and clinical facilities. It has also the potential to reduce the experimental errors associated with separate irradiations.

**Figure 2.**
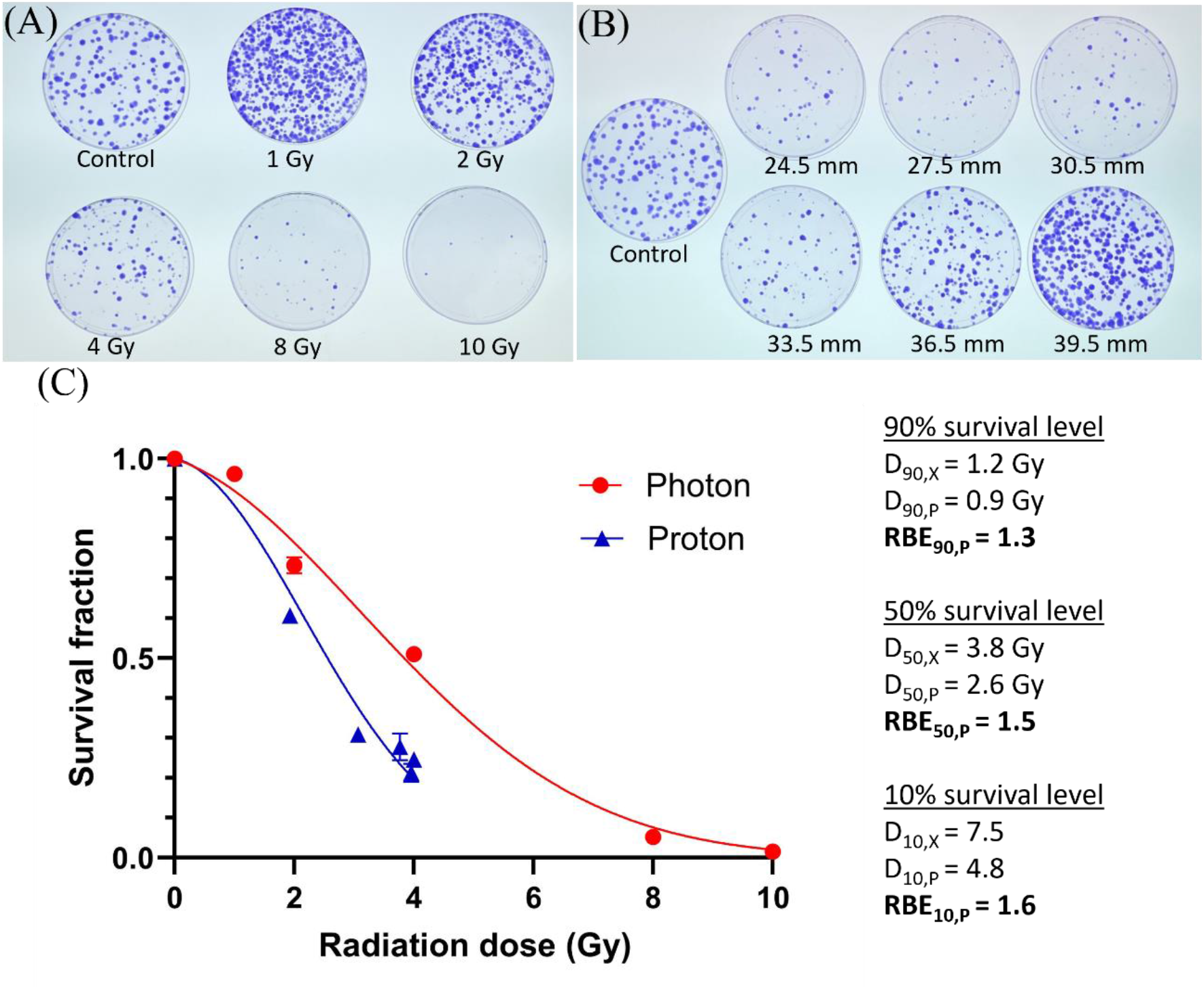
Clonogenic survival assay for U87 cells irradiated with photon (A) and proton (B); (C) Cellular survival fraction as a function of radiation dose and linear quadratic model fitting.

### Determination of RBE based on flow cytometry analysis of DNA DSBs repairs

Next, we used the microtissue cassette to measure the RBE based on the standard γ-H2AX assay used to assess DSBs repairs. The fluorescence intensities as measured by flow cytometry for γ-H2AX foci versus position at distance from the center of the SOBP for 1 Gy proton irradiation is shown in Fig. 3A. At 1h following post-irradiation, the fluorescence signal of γ-H2AX foci was found to correlate well the percentage of the deposition dose. In parallel, three microtissue slides were placed in one cassette and irradiated with photon irradiation at 1 Gy. The U87 cells on these slides were fixed at 1h and analyzed with flow cytometry. Of note, microtissue samples for both proton and photon irradiation were prepared, irradiated and analyzed on the same day to reduce the bias as much as possible. The RBEs for the six positions within the SOBP and at the distal fall-off were calculated following the methodology previously reported for the γ-H2AX assay.^6^ As shown in Fig. 3B, proton RBEs for 1 Gy were calculated to be 1.12 ∼ 1.33 within the SOBP (positions A, B, C in Fig. 1F) but increased at the distal fall-off to 1.58 (position D) and 1.38 (position F). These RBEs are consistent those determined with the U87 cell survival fraction clonogenic assay (RBE = 1.3∼1.6) shown in Fig. 2C. In addition, the physical dose, corrected dose (assuming a uniform RBE of 1.1) and biological dose (physical dose times calculated RBE) are plotted to demonstrate the correlation between the RBE and the biological dose as shown in Fig. 3C. The result shows a higher effective biological dose at all points along the SOBP and distal fall-off. There are some limitations associated to flow cytometric measurements of the fluorescent signals compared to microscopy image-based enumeration of foci associated to γ-H2AX foci to determine the RBE as the high background noise limits the accuracy of the test for low foci numbers.

**Figure 3.**
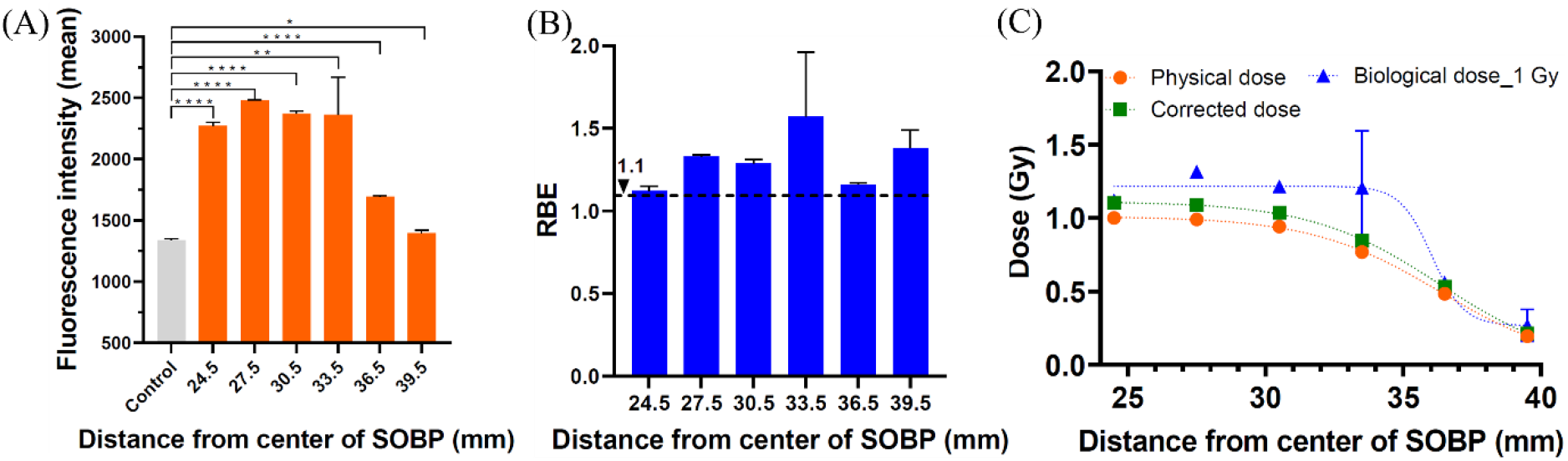
(A) γ-H2AX fluorescence intensities and (B) calculated RBEs as a function of distance from the center of the SOBP for six tested positions; (C) Plotting of the physical dose, corrected dose (physical dose*1.1) and biological dose (physical dose*calculated RBEs) for 1 Gy proton dose.

### Foci analysis along SOBP

Radiation treatment triggers DNA double strand breaks in the nucleus in the first place, followed by a cascade of cellular pathways which may eventually lead to cell death. It has been proposed that particle radiation therapy including proton and carbon ions causes complex DNA damage com-pared to x-ray. To further explore the utility of the 3D printed microtissue cassette in measuring and quantifying γ-H2AX foci, we conducted a detailed analysis of the foci numbers and their spatial arrangement at various positions in the SOBP and distal fall off. γ-H2AX foci were analyzed 1 h after 1 Gy of proton irradiation (Fig. 4). The number of foci was steady at 21-24 per cell and decreased to < 15 in the distal fall-off region, indicating a good correlation with the PDD. Track-like spatial arrangements of foci were not observed in the post Bragg peak region unlike in a previous report.^8^ This difference might be because cells were positioned in our set up perpendicular to the direction of the proton beam. It has also been shown that Bragg-peak protons induce larger and more irregularly shaped γ-H2AX foci which may in-crease the complexity of DNA repairs.^9^ As shown in Fig. 4H, the size of the γ-H2AX foci did not correlate with the proton dose along the SOBP and distal fall-off although a statistically non-significant difference was observed at the most distal position (39.5 mm) which may be attributed to the low proton dose (0.195 Gy).

**Figure 4.**
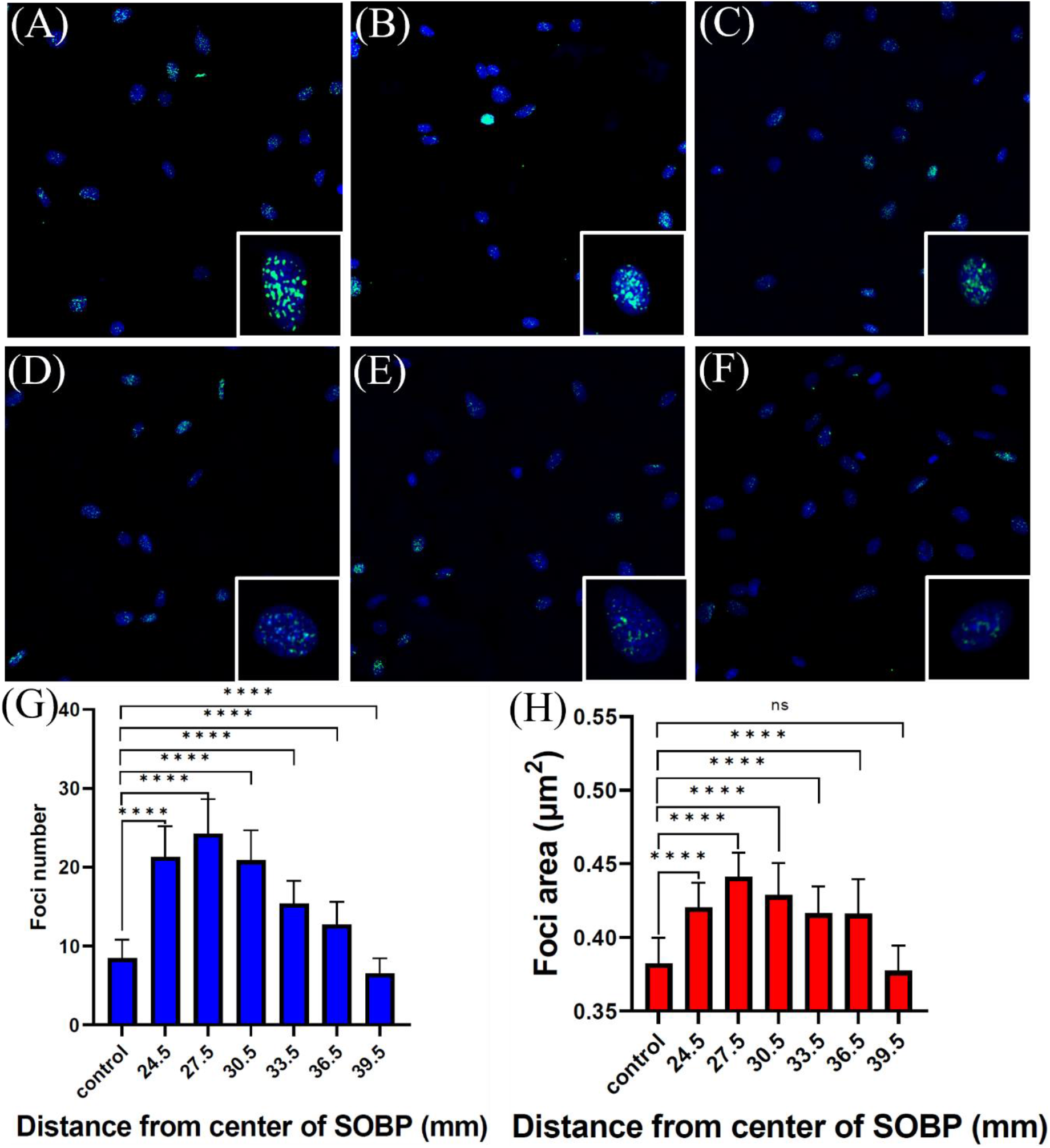
High content imaging of γ-H2AX foci in U87 cells irradiated with 1 Gy proton at distances from the center of the SOBP of (A) 24.5 mm; (B) 27.5 mm; (C) 30.5 mm; (D) 33.5 mm; (E) 36.5 mm; (F) 39.5 mm; (G) Foci numbers and (H) foci areas as a function of distance from the center of SOBP calculated from random 25 areas (n = 3) for each SOBP position.

## Conclusion

We report on an innovative and reliable experimental approach based on 3D printed microtissue cassettes to deter-mine the proton RBE along the SOBP including in the distal fall-off using clonogenic assay and *in vitro* γ-H2AX assay. This design enabled high spatial resolution and high throughput measurements, two important features for such radiobiological measurements in clinical radiotherapy facilities. The presented results provide further evidence of the underestimation of the proton RBE at SOBP and distal fall-off region and contribute to address the current ambiguity. More generally, we envisage that this approach provides a simple and universal method to facilitate addressing critical radiobiological questions associated to charged particle-based radiation therapy including helium and carbon ions.

## ACKNOWLEDGMENT

Dr Chih-Tsung Yang would like to thank the funding support from The Hospital Research Foundation Group (C-F-EMCR-008). This work used the NCRIS and Government of South Australia enabled Australian National Fabrication Facility - South Australian Node (ANFF-SA). We thank the Radiation Research Core Laboratory at Linkou Chang Gung Memorial Hospital for their assistance.

